# Label free, capillary-scale blood flow mapping *in vivo* reveals that low intensity focused ultrasound evokes persistent dilation in cortical microvasculature

**DOI:** 10.1101/2024.02.08.579513

**Authors:** YuBing Shen, Jyoti V. Jethe, Ashlan P. Reid, Jacob Hehir, Marcello Magri Amaral, Chao Ren, Senyue Hao, Chao Zhou, Jonathan A. N. Fisher

## Abstract

Non-invasive, low intensity focused ultrasound (FUS) is an emerging neuromodulation technique that offers the potential for precision, personalized therapy. An increasing body of research has identified mechanosensitive ion channels that can be modulated by FUS and support acute electrical activity in neurons. However, neuromodulatory effects that persist from hours to days have also been reported. The brain’s ability to provide targeted blood flow to electrically active regions involve a multitude of non-neuronal cell types and signaling pathways in the cerebral vasculature; an open question is whether persistent effects can be attributed, at least partly, to vascular mechanisms. Using a novel *in vivo* optical approach, we found that microvascular responses, unlike larger vessels which prior investigations have explored, exhibit persistent dilation following sonication without the use of microbubbles. This finding and approach offers a heretofore unseen aspect of the effects of FUS *in vivo* and indicate that concurrent changes in neurovascular function may partially underly persistent neuromodulatory effects.

## Introduction

Low-intensity focused ultrasound (FUS) is an emerging technology for non-invasive neuromodulation. Compared with other non-invasive modalities such as transcranial magnetic stimulation (TMS) or transcranial direct current stimulation (tDCS), FUS offers the potential for restricting modulatory stimuli to within volumes that are ∼3-5 orders of magnitude smaller and with millimeter-scale cross-sectional focal area ^1–5^. These resolution advantages offer the possibility for minimizing off-target effects as well as the potential for more precise, personalized treatment. In human subjects, FUS has been explored as a therapeutic for a wide range of conditions such as stroke ^6^, Alzheimer’s disease ^7,8^, essential tremor ^9,10^, Parkinson’s disease ^11,12^, as well as chronic pain ^13^. Beyond therapy, FUS has been shown to acutely modify sensory evoked neural responses in small animal models ^14^, swine ^15^, nonhuman primates ^16^, and humans ^17^. Given that FUS has the capability of evoking sensations including somatosensory ^18–22^ and visual ^23,24^, this modality has presented new stimulus methods for brain-computer-interface applications ^25–27^.

Central to wider clinical adoption is a clearer understanding of dosage and effects. The parameter space for designing stimuli and dosing is wide, and altering individual parameters such as duty cycle has been found to qualitatively change the effects on the brain and their associated timescales^28^. For example, in some instances, altering duty cycle was found to shift effects from excitatory to inhibitory even when the average pressure is not changed ^29^. Recent work on mechanisms has provided compelling evidence that FUS elicits action potentials in neurons by activating mechanosensitive Ca^2+^ channels that include TRPP1, TRPP2, TRPC1, and Piezo1 ^30^. FUS stimulation has also been shown to increase Ca^2+^ influx in astrocytes by opening TRPA1 channels, resulting in glutamate release through Bestrophin-1 (Best1) channels ^31^.

Beyond acute electrophysiological effects, FUS can elicit persistent plasticity ^2,16,32^ up to weeks, in some studies ^33,34^. In terms of underlying mechanisms for these chronic effects, FUS appears to modulate the function of NMDA receptors ^35^, a key contributor to many forms of synaptic plasticity ^36–38^. Zhao et al. (2023) suggested that FUS-induced increases in expression of brain-derived neurotrophic factor (BDNF), glial cell line-derived neurotrophic factor (GDNF), and vascular endothelial growth factor (VEGF) could underly the increased growth rate of dendritic spines that they observed over a 28-day course of repeated doses of FUS ^8,39^.

Sonication-evoked cerebrovascular effects and hemodynamics, generally considered secondary to neuronal activity ^40^, could also underly at least some of the observed neuromodulatory effects. Neuronal activity elicits cerebral blood flow and cerebral blood volume changes that are spatially restricted to roughly the same areas that are electrically active. This phenomenon, termed neurovascular coupling, is the result of an intricate choreography of multiple cell types and signaling mechanisms ^41^ and perturbing vasculature would impact neuronal function. In fact, pathological perturbation of even single capillaries can cause larger-scale changes in blood flow ^42^ and may result in microvascular ischemia ^43^. In terms of potential mechanisms, some of the same mechanosensitive channels found to be critical for FUS-induced neural effects are also expressed in vasculature. TRPC1, for example, is expressed in vascular endothelial cells ^44–46^, TRPP1 in vascular smooth muscle cells ^47–49^, and Piezo1 is expressed in both cell types and also plays a critical role in vascular development and remodeling ^50–52^. Because hemodynamic responses lag neural activation, vascular effects could potentially contribute to persistent apparent neuroplasticity.

The impact of FUS on cerebral hemodynamics has been studied using invasive and noninvasive methods. Functional magnetic resonance imaging (fMRI), for example, has been instrumental in noninvasively exploring the deep brain effects of FUS neuromodulation as well as its viability in larger brains including humans. fMRI has revealed FUS-induced hyperaemia both cortically and subcortically by means of volumetric blood-oxygen-level-dependent (BOLD) signals ^16,22^. Near infrared spectroscopy (NIRS) ^53^, another non-invasive approach for monitoring cerebral hemodynamics, has been used to measure hemodynamic signals evoked by administering FUS to the somatosensory cortex ^54^. While generally lacking the spatial resolution of fMRI, NIRS can monitor hemodynamics with much higher temporal resolution^55^. Kim et al. found that the FUS-induced hemodynamic response, measured in terms of inferred changes in oxy- and deoxyhemoglobin concentrations, largely resolves within 20 seconds. That timescale is similar to sensory-evoked hyperaemia measured using NIRS^54^. Unlike noninvasive measurements, which have relatively coarse spatial resolution, invasive imaging techniques such as laser speckle contrast imaging ^56,57^ and optical intrinsic signal imaging (OISI; Yuan et al., 2023) have resolved FUS-induced flow changes and dilation in individual vessels. Mirroring the NIRS findings, these studies have similarly observed FUS-induced hemodynamic responses in individual vessels largely subside within ∼10 seconds.

While the results of FUS-associated hemodynamic measurements seemingly suggest that acute effects are likely of neuronal or astrocytic origin and that persistent neuromodulatory effects do not manifest in prolonged vasomodulation, none of these experiments has heretofore directly resolved the impact of FUS alone on microvasculature. For widefield optical techniques such as laser speckle and OISI, this is partly owing to relatively poor axial resolution due to the use of low numerical aperture lenses. Measuring the functional response in small vessels is critical because microvascular branches ∼3−15 μm in diameter, which include capillaries, small arterioles, and venules, are at the frontline for neurovascular coupling ^41,59^ and also likely easier to perturb mechanically by FUS ^60^.

To obtain insight into a potential microvascular reflection of persistent effects following sonication, we developed a label-free optical approach for interrogating dilation in vasculature down to capillary scale *in vivo* directly at the site of ultrasound delivery. Our apparatus used a custom spectral domain optical coherence tomography angiography (OCT-A) device that resolves vasculature based on light scattering changes due to moving red blood cells. Compared with other optical modalities for resolving blood flow at microvascular scales, OCT-A offers the advantage of being label free, in contrast with other optical techniques such as two-photon microscopy, which requires intravenous tracers or targeted expression of fluorescent reporters. Additionally, because the depth resolution in OCT is limited by the characteristics of the broadband light source rather than an objective’s optical depth of field ^61^, relatively low-numerical aperture, long working distance lenses can be used to survey large brain regions without sacrificing depth resolution. Integrating this with a wide-aperture ring transducer whose acoustic and optical foci were co-localized, we were able to obtain a direct glance into the effects of FUS on cortical microvasculature.

## Results

### In situ, label-free microvascular-level measurements of dilation during FUS neuromodulation

Using a custom spectral domain OCT-A apparatus (Fig. 1A) that was able to compensate for long working distances that traversed ∼10 cm of water within the ring transducer, we were able to obtain 3D portraits of vasculature (Fig. 1B) with 2.48 × 3.48 μm (axial × lateral) resolution at depths upwards of 650 μm, label free in wild-type mice. Whereas techniques such as MR-acoustic radiation force imaging (MR-ARFI) can be used to localize the FUS focus in fMRI experiments ^62^, a challenge for targeting FUS in epi-illumination optical systems *in vivo* is that the FUS beam is not intrinsically optically detectable. Because small positioning errors can lead to uneven sonication patterns within an optical field of view, we used a phased array ring transducer with a beam width larger than the analyzed zones (Fig. 1C) to ensure that our analysis was restricted to vessels that received the same FUS dosing. Additionally, as depicted in Fig. S5, the axial profile of the beam reveals an extruded focus whose intensity profile had a full-width-half-max (FWHM) of ∼5 mm. The sonication focal profile would be minimally altered by tissue and therefore can be considered relatively uniform throughout the imaging depth.

**Figure 1:**
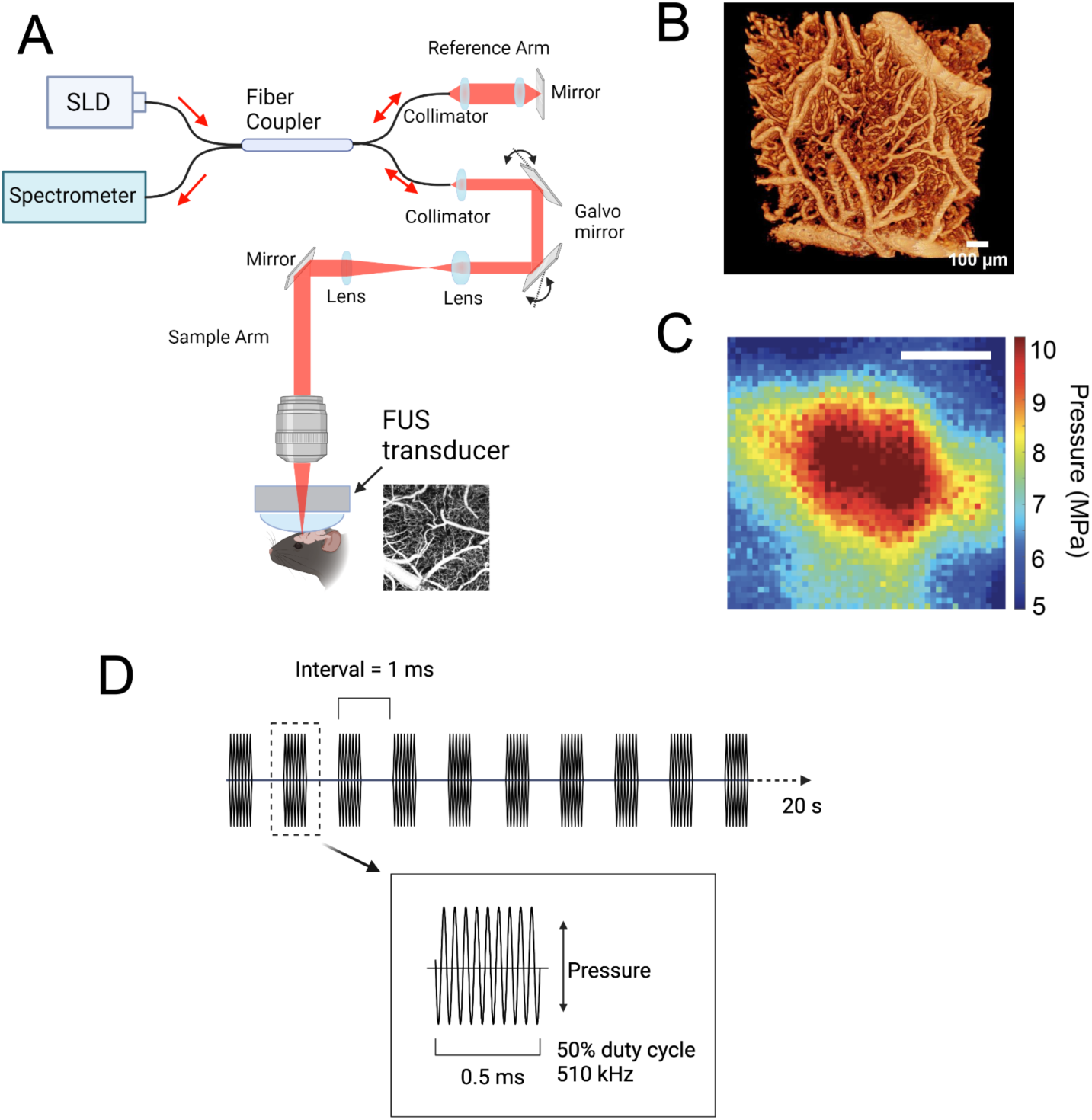
Combined optical coherence tomography angiography (OCT-A) and FUS for observing multi-scale vascular responses to sonication. **A** depicts a schematic for a custom frequency domain OCT-A apparatus with in-line FUS ring transducer. The inset shows a sample 1 mm × 1 mm *en face z*-projection revealing flow patterns measured in somatosensory cortex. **B** shows a sample 3D surface mesh rendering of vascular architecture, as reconstructed from OCT-A imagery. **C** shows the lateral profile of the FUS beam (scale bar = 1 mm), and **D** illustrates the pulsed sonication protocol; sonication was delivered as a 20 s period of 510 kHz ultrasound pulsed at 1 kHz with 50% duty cycle.

Compared with vascular analysis and tabulation based on two-photon imaging^63^, the vessel morphology is of lower signal-to-noise. We therefore developed an image processing pipeline that was able to selectively segment vessel branches based on diameter, thereby permitting us to selectively analyze dilation dynamics in diverse vessel populations. Fig. 2 shows *z*-projections taken over narrow ranges centered at the cortical surface and ∼340 μm below, illustrating that there is greater representation of small vessels at these deeper depths. This can be visualized more quantitatively in Fig. 4A, which shows the distribution of vessel diameters, binned by depth (*N* = 10 animals). A color-coded *z*-projection showing the full depth span of angiography data is shown in Fig. 2A. A visualization of the vessel segmentation process is depicted in Fig. S3.

**Figure 2:**
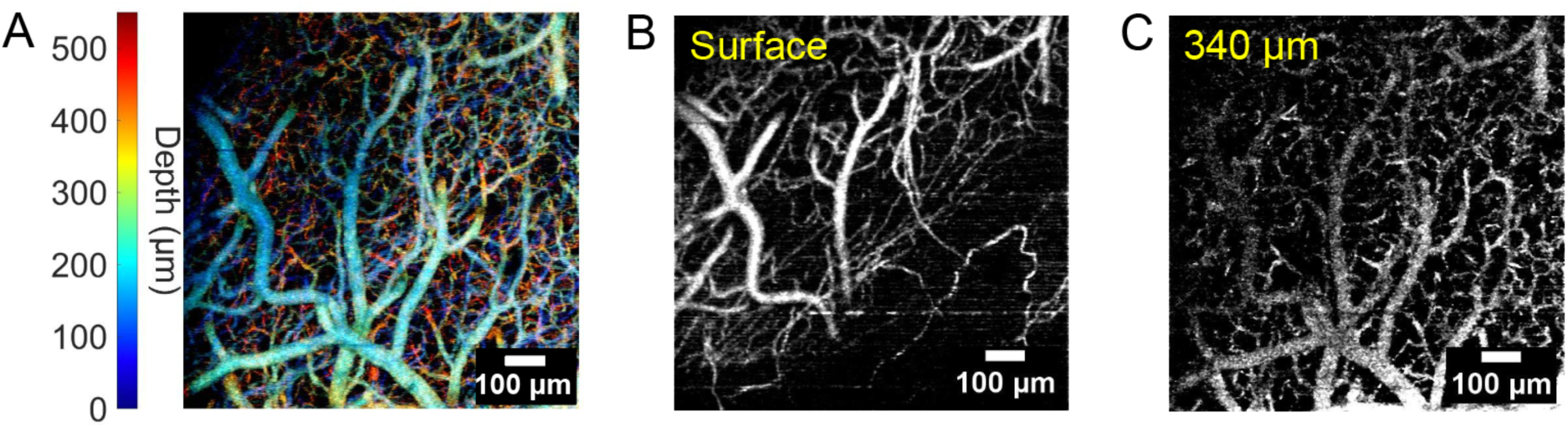
Depth-resolved flow patterns in the mouse somatosensory cortex. **A** shows data from all depths, which are here encoded by color. As is apparent, consistent with known cerebrovascular architecture, the deepest features resolve a fine mesh of narrow vessel branches. **B** and **C** show *z*-projections from narrow ranges of lateral slices centered at the cortical surface and ∼340 μm below. The depth range for projections in **B** and **C** is ± ∼2 μm axially.

### FUS elicits vascular dilation most prominently in small-diameter vessel branches

Averaged over sonicated animals, excluding those used strictly for immunohistochemical analysis, sonication at 510 kHz FUS (50% duty cycle, and 1 kHz pulse repetition) delivered at spatial-peak pulse-average intensities (*I_sppa_*) of 1 W/cm^2^ (*N* = 10) and 10 W/cm^2^ (separate *N* = 10) for 20 seconds elicited dilation that was significant for the population of vessels of diameter less than 15 μm, but not for populations of larger diameter vessels (Fig. 3 & Table 1). Prior to FUS, the average diameter of the small vessel population was 6.2 ± 1.3 μm (mean ± standard deviation); within 1 minute following 1 W/cm^2^ sonication, this vessel population exhibited an 17% ± 3% increase in diameter (*P* < 0.0001, using a two-tailed paired Student’s t-test). Sonicating at a higher intensity (10 W/cm^2^) yielded a greater dilation of 27% ± 8% (*P* < 0.0001), which was significantly different from the result of 1 W/cm^2^ (*P* = 0.0052 via one-way ANOVA and Fisher’s Least Significant Difference test). At both intensities, dilation in small vessels following an initial sonication persisted beyond 10 minutes but did not significantly change in magnitude (Fig. S1).

**Figure 3:**
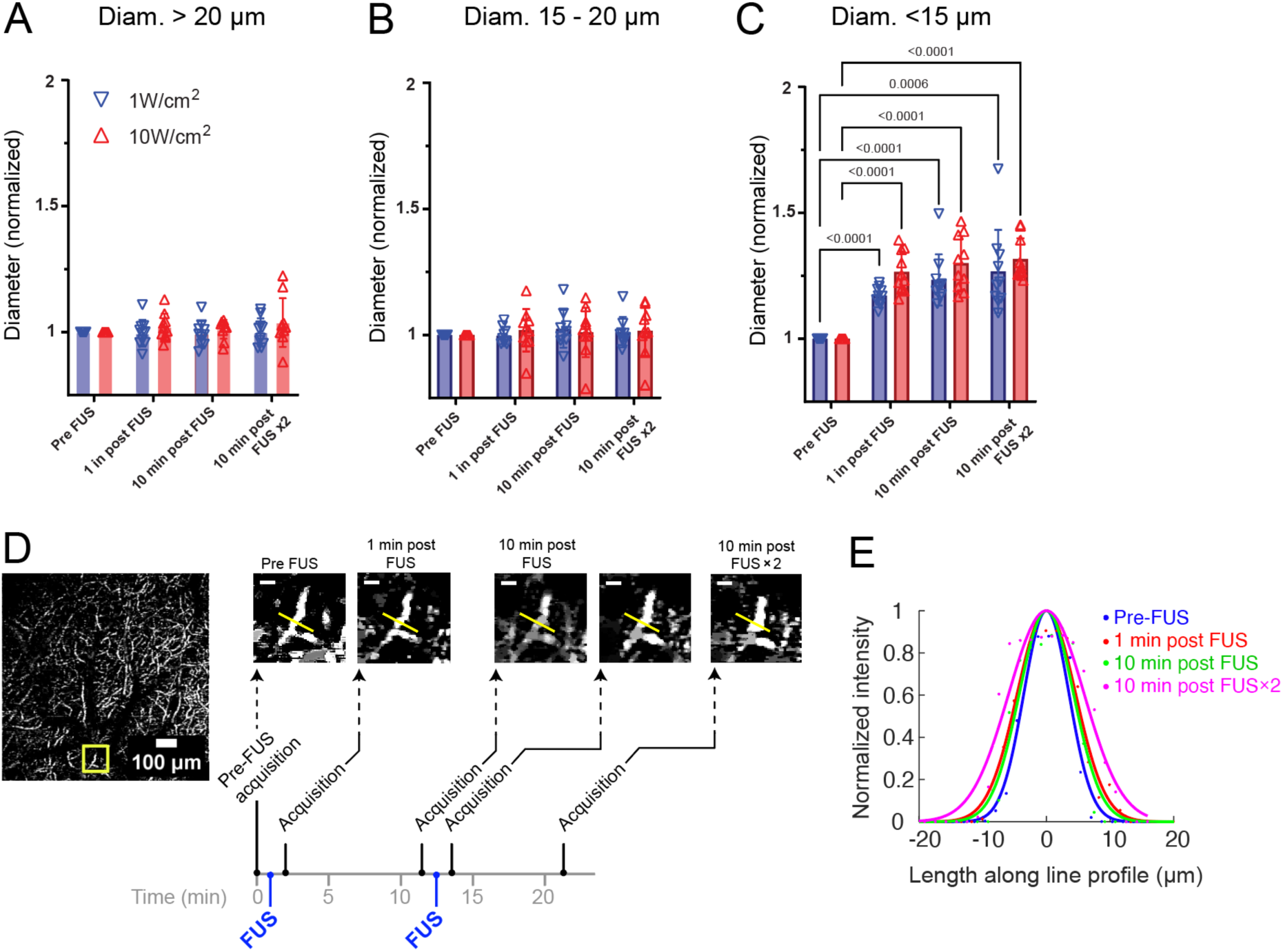
Dissecting vascular effects of FUS *in vivo*. Normalized changes in vessel branch diameters, averaged over all animals, are binned based on pre-FUS diameter ranges: >20 μm (**A**), 15-20 μm (**B**), and <15 μm (**C**). Vessels of pre-FUS diameter less than 15 μm, but not larger vessels, displayed significant increases in diameter that persisted beyond 10 minutes. Values displayed atop brackets in **C** report *P*-values that were below 0.05, obtained using a paired, two-tailed Student’s t-test. Individual data points correspond to results from individual animals. **D** illustrates the sonication and imaging protocol in the form of a timeline and shows OCT-A monitoring in a single vessel branch during the course of consecutive doses of FUS. The yellow square in the left-most image shows a region that is expanded in images to the right. The dimensions for the full image on left are 1 mm × 1 mm, and the dimensions image for smaller images on the right are 110 μm × 110 μm (scale bar: 20 μm). Sonication consisted of pulse bursts that were delivered at a rate of 1 kHz over a period of 20 sec. **E** shows Gaussian fits to the intensity profiles measured across the yellow lines shown in the smaller image panels of **D**; the dots show actual intensity values, and are derived from the one representative animal whose images are shown in **D**. The intensity profiles were normalized to visualize changes in diameter, which can be approximated by the full-width-half max of the Gaussian fits.

**Table 1:** Analysis results from FUS treatment averaged among all experiments. Values following treatment are reported as normalized diameters. The FUS × 1 and FUS × 2 refer to the first and second sonication doses (see timeline in Fig. 3D). In addition to the changes summed over all vessel branch diameters, analysis is reported for selected diameter ranges. *P*-values were obtained using a paired, two-tailed Student’s t-test that compared individual animals. Diameter values are reported as mean ± standard deviation.

| Diam. range | FUS intensity<br>($I_{\text{sppa}}$) | Diameter<br>pre-FUS $\times 1$<br>($\mu\text{m}$) | Normalized<br>diameter, 1 min<br>post FUS $\times 1$ | Normalized<br>diameter, 10<br>min post FUS | Normalized<br>diameter, 10 min<br>post FUS $\times 2$ | *P*-value of post FUS<br>$\times 1$ comparison with<br>pre-FUS |
| --- | --- | --- | --- | --- | --- | --- |
| All vessels | Sham (n = 3) | 18.70±19.79 | 1.00±1.01 | 1.08±1.05 | 1.04±1.08 | - |
| All vessels | 1 W/cm<sup>2</sup> (n = 10) | 16.27±16.01 | 1.05±0.92 | 1.06±0.91 | 1.06±0.93 | 0.28 |
| All vessels | 10 W/cm<sup>2</sup> (n = 10) | 19.40±14.39 | 1.02±0.73 | 1.01±0.71 | 1.01±0.69 | 0.77 |
| > 20 $\mu\text{m}$ | Sham | 38.94±23.27 | 0.98±0.58 | 0.96±0.58 | 0.98±0.60 | - |
| > 20 $\mu\text{m}$ | 1 W/cm<sup>2</sup> | 35.25±19.16 | 1.00±0.05 | 0.99±0.05 | 1.00±0.05 | 0.64 |
| > 20 $\mu\text{m}$ | 10 W/cm<sup>2</sup> | 31.63±13.43 | 1.02±0.05 | 1.01±0.04 | 1.04±0.09 | 0.87 |
| 15 - 20 $\mu\text{m}$ | Sham | 17.28±1.35 | 1.01±0.08 | 0.97±0.06 | 0.99±0.06 | - |
| 15 - 20 $\mu\text{m}$ | 1 W/cm<sup>2</sup> | 17.66±1.38 | 1.00±0.03 | 1.02±0.07 | 1.01±0.06 | 0.6 |
| 15 - 20 $\mu\text{m}$ | 10 W/cm<sup>2</sup> | 17.57±1.40 | 1.02±0.08 | 1.01±0.09 | 1.02±0.09 | 0.45 |
| < 15 $\mu\text{m}$ | Sham | 6.50±1.58 | 0.99±0.01 | 1.00±0.01 | 1.00±0.004 | - |
| < 15 $\mu\text{m}$ | 1 W/cm<sup>2</sup> | 6.20±1.27 | 1.17±0.03 | 1.23±0.10 | 1.27±0.16 | **8.6E-26** |
| < 15 $\mu\text{m}$ | 10 W/cm<sup>2</sup> | 7.48±2.24 | 1.27±0.08 | 1.30±0.1 | 1.32±0.08 | **3.6E-08** |

As depicted in Fig. 3A-B, for vessel branches of larger diameters, FUS evoked small changes in dilation that were non-significant. Vessel segmentation and analysis at the microvascular level also enabled us to catalog and track individual vessel branches; Figs. 3D-E follow the dilation of one tracked sample vessel branch following sonication. In sham experiments (*N* = 3), vessel branches, regardless of diameter range, did not exhibit significant changes at any time point (Table 1).

### Depth-dependent dilation effects of FUS

FUS-induced effects differed not only based on a vessel branch’s baseline diameter, but also based on depth. Fig. 4A displays the distribution of vessel branch diameters as function of depth prior to sonication in the cohort of animals subsequently sonicated at 10 W/cm^2^. While there were a range of diameters at each depth (binned in 25-μm increments), the mean branch diameter decreased in depth progressively from 125 μm toward 375 μm. This profile of depth-dependence in vessel diameter is consistent with prior results from two-photon microscopy (e.g. Blinder et al., 2013)^63^.

**Figure 4:**
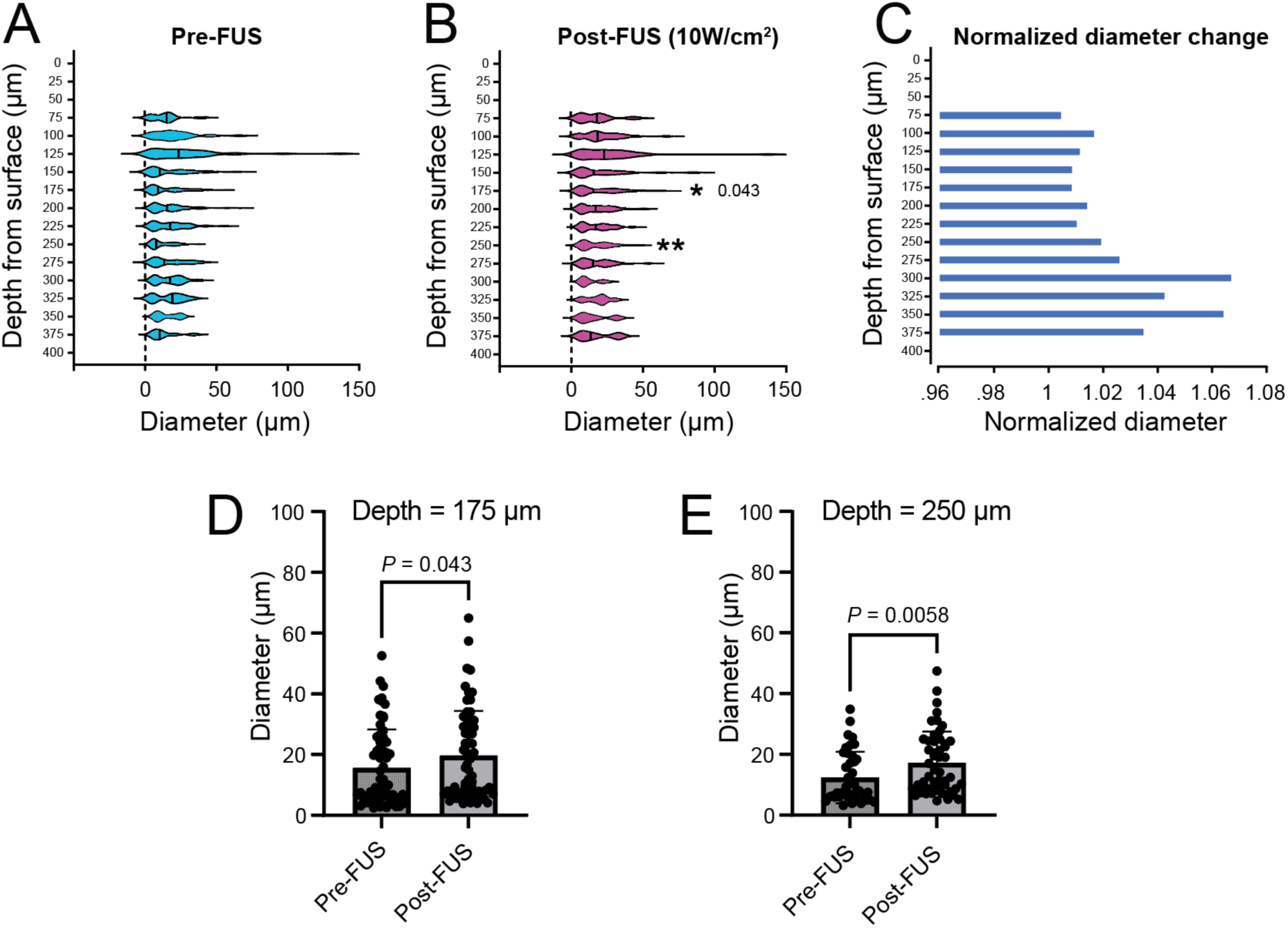
Depth-resolved diameter analysis of FUS-induced vascular dilation. **A** shows the distribution of vessel diameters binned in depth from the surface of the cortex down to 375 μm. Each depth measurement is the result of binning in units of 25 μm. Results at each depth are presented as violin plots and represent all vessels at that depth bin and are derived from the results of 10 animals that were exposed to 10W/cm^2^ FUS. Small vertical black lines within each violin plot represent the average value. **B** shows the depth-dependence of vessel branch diameters from those same animals after treatment with 10W/cm^2^ FUS. Significance was measured using a Mann-Whitney-Wilcoxon test. **C** depicts normalized post-FUS changes in average vessel diameter with reference to baseline values in **A**. **D** and **E** show bar charts that feature actual datapoints, each of which corresponds to an individual vessel branch, within the depth range centered at 175 μm and 250 μm respectively.

This analysis also enabled us to identify depth-dependent changes in vessel branch diameter upon treatment with FUS. As depicted in Fig. 4C, the largest mean changes in diameter were at depths below 225 μm. Comparing individual depth bins (Fig. 4D-E), significant differences were found at depths of 175 μm and 250 μm upon sonication with 10 W/cm^2^ (*P* = 0.043 and 0.0058, respectively using a Mann-Whitney-Wilcoxon test).

### Impact of FUS on vascular permeability and tissue integrity

At high intensities, FUS can permeabilize the blood-brain barrier (BBB) and impact neurovascular response^64^. To address the possibility that the vascular responses we observed were due to tissue damage or extravasation, in a separate cohort of animals (*N* = 5) we assessed vessel morphology and leakage of intravenously-injected Evans blue dye, which binds to serum albumin and does not leave the vasculature unless the BBB is permeated. We exposed four of these mice (the remaining animal was used as an unsonicated sham) to FUS within the intensity range used in these experiments (1.2 W/cm^2^) and, as a positive control for tissue damage, high intensity (200 W/cm^2^), at adjacent locations on the same cortical hemisphere (Figs. 5 and S4). After 24 hours, sites exposed to high-intensity FUS displayed clear tissue damage as well as microvascular disruption, as visualized by vascular marker CD31. Regions targeted with high intensity FUS also displayed extravasation of Evans blue dye in the parenchyma. These effects extended beyond 350 μm in depth. In contrast, exposure to a low intensity within the range of the other experiments (1.2 W/cm^2^) yielded no evidence of gross tissue disruption nor Evans Blue dye leakage.

**Figure 5:**
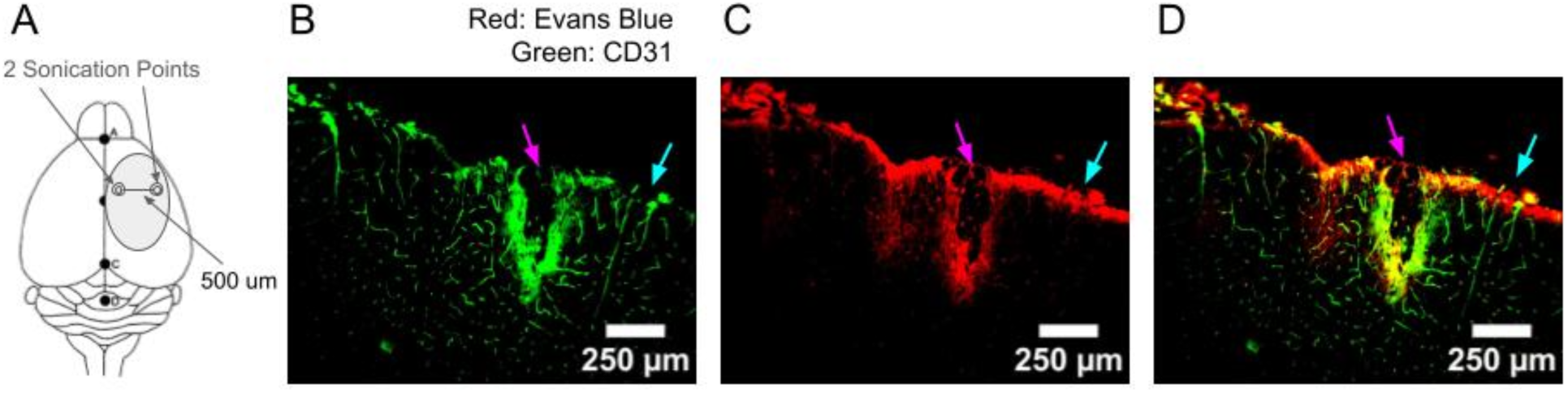
Immunohistochemical assessment of vascular integrity and BBB permeabilization. **A** shows an illustration of the relative coordinates on the brain where FUS was targeted. The medial location received higher intensity sonication (>200 W/cm^2^ *I*_sppa_, 100% duty cycle, 20 s), while the lateral location was sonicated at a lower intensity (1.2 W/cm^2^, 100% duty cycle for 20s). **B-D** show coronal sections obtained at the rostrocaudal axis location where sonication was performed. The magenta and cyan arrows represent, respectively, high-and low-intensity sonication locations. CD31 labeling is displayed in **B**, Evans blue in **C**, and **D** shows a merged image. Results of an additional 3 animals and sham experiment are shown in Fig. S4.

## Discussion

The results we obtained fill a gap in exploration of vascular responses to FUS, which have heretofore measured either responses in larger vessels or in diffuse, bulk tissue hemodynamics. The closest relevant studies involved the use of two-photon microscopy to dissect microvascular changes following the combination of FUS and microbubbles^60,65,66^. Because microbubbles significantly amplify mechanical displacement due to volume oscillations, the underlying mechanisms in findings from that body of research are likely different. Those studies largely report vasoconstriction, though Cho et al. also observed modest vasodilation in small diameter vessel branches. The authors suggested that these effects could potentially be related to secondary effects of BBB-disruption, namely inflammatory responses ^67^. This mechanism may not be unique to FUS; in the case of tDCS, for example, despite the fact that the modulatory stimulus is electrical current rather than mechanical perturbation, preclinical studies have revealed evidence that persistent plasticity may be similarly associated with an inflammatory vascular response due to perturbed BBB permeability ^68^. Although our observed vasoconstrictive trend in vessel branches larger than 20 μm in diameter were not statistically significant, that combined with vasodilation of smaller diameter vessels bears semblance to those previous results. Additionally, because individual vessels vary in their diameters along their length, particularly at branch points for larger vessels, quantification of potential vasoconstrictive effects could have been diluted given that microvessels represented the majority of identified vessels. At the same time, opposite trends—FUS-induced dilation of large vessels—have been observed using laser speckle imaging without microbubbles^56^. Previous microvascular observations may therefore indeed be of limited relevance.

Despite the absence of microbubbles and the fact that we found no evidence of Evans blue dye leakage, it is not unfathomable that mechanical perturbation could have activated an inflammatory response due to a small amount of BBB permeabilization that our immunohistochemical analysis was not able to resolve. Reports that have measured Evans blue leakage with other methods such as near-infrared whole brain imaging, however, have also observed persistent neuromodulation in the absence of BBB permeabilization evidence ^69^. This provides some supporting evidence that FUS alone, without BBB permeabilization can lead to persistent effects. To that point, it should be noted that *in vitro* results provide evidence for entirely vascular-independent components of persistent effects; Clennell et al. (2021), for example, found that FUS can induce up to 8 hours of sustained enhanced excitability in cultured cortical neurons, possibly due to dephosphorylation of select subunits of voltage-gated potassium channels (i.e. K_v_2.1) ^71^.

The fact that the dilation we observed in microvasculature was persistent and not associated with sensory activation suggests that the effect is not simply a reflection of FUS-triggered hyperaemia secondary to acute neural activation; for instance, we have previously shown that acute, sensory activation evokes increases in capillary blood flow that subside within ∼10 sec ^72^. If our microvascular trends do indeed reflect a response to neuronal effects, they may be more likely related to the prolonged elevated excitability described by Clennell et al. (2011).

It is likely that the prolonged dilation is related to a combination of direct and indirect effects of FUS modulation of mechanosensitive channels in all key cell types in the neurovascular unit, i.e. astrocytes, endothelial cells, smooth muscle cells, pericytes, and of course neurons. As noted previously, endothelial cells and smooth muscle cells express some of the same FUS-sensitive mechanosensitive channels as in neurons, all of which have recently been found to be directly sensitive to FUS, at least in culture ^73^. We would include in “indirect” effects the FUS-dependent release or effect of vasodilators such as nitric oxide (NO). This could originate from dilation in larger, smooth muscle cell-lined vessels in response to either FUS-induced increased NO release from astrocytes as described by Oh et al. ^31^, or else enhancement of NO synthase, as suggested by Yuan et al. (2018). At the capillary level, the increase in NO concentration could cause dilation by inducing relaxation of pericytes ^74^.

While our experimental methods did not permit us to directly explore the involvement of specific cell types, the depth-resolved analysis of our imaging results may offer indirect supporting evidence for the role of pericytes. The diameter of a vessel branch in the cortical microvasculature is inversely related to the “branch order,” i.e. the number of bifurcations with respect to an individual penetrating arteriole that have occurred leading to the branch^75^. Branches of higher order feature sequentially different cell types; Grant et al., for example, observed that branch order in the mouse cortical vasculature predicts both the presence and type of pericytes found^76^. Based on the vessel diameter dependence that we observed associated with FUS-induced effects, our findings suggest a strong bias toward branches of higher order. Unfortunately, resolving features in depth such as branch order in OCT-A imaging is complicated by projection artifacts, or “tailing” underneath vessels which is caused by multiple-scattering^77^. These artifacts can be potentially mitigated using computational approaches such as convolutional neural network deep-learning techniques^78^. While this analysis is outside of the scope of the current report, future work will certainly pursue this line of investigation.

Prior work that used two-photon imaging to identify vessel branching patterns, however, may offer an alternative approach to estimating branch order. In particular, Blinder et al. found that arteriolar branching probability peaked at deeper layers (roughly depths beyond 200 μm) than venules, which peak within the first ∼100 μm from the cortical surface^63^. In the context of this previous work, the diameter and depth biases that we observed suggest, albeit indirectly, that FUS-induced dilation occurs most prominently in high branch order pre-capillary arterioles and capillaries.

The observation that dilation changes are potentially most prominent for vessel branches that feature pericytes rather than smooth muscle cells could be significant given the longstanding controversy regarding the role of pericytes in regulating neurovascular coupling^79^. Recent work on this topic has found that pericytes are at least sufficient for maintaining basal capillary tone, if not facilitating dilation^80^. Notably, pericytes have been found to be significantly slower than smooth muscle cells in their mechanical response ^81^ which is correlated with their lower expression of ɑ-smooth muscle actin (ɑ-SMA). Given their inherently slower kinetics, if the identity (e.g. smooth muscle cell devoid vessel branches) of the vessel population we observed to have the greatest change could be confirmed, this could suggest that dysfunction in pericytes, which have inherently slower kinetics than smooth muscle cells, is an important contributor to slow, persistent neuromodulatory effects.

The use of anesthesia presents potential confounds for this and the majority of other FUS neuromodulation studies in small animals. On their own, many anesthetics alter hemodynamics and vascular function owing to a wide range of mechanisms including altered cerebral autoregulation, vasomotor reactivity, and neurovascular coupling ^82^. Yoo et al. ^83^ found that FUS administered to the thalamus could alter anesthesia state, specifically for ketamine-xylazine. FUS administered to brainstem nuclei, particularly the dorsal raphe nucleus, has also been shown to accelerate recovery from isoflurane-induced anesthesia ^84^. Decades of experimental and theoretical research has shown that systems-level thalamocortical oscillations and their interplay with brainstem nuclei serve a controlling role in regulating consciousness under anesthesia ^85–93^. It is unsurprising, therefore, that directing FUS subcortically to those regions affects recovery from anesthesia. In our experiments, FUS was directed laterally at the somatosensory cortex, intentionally minimizing the possibility of affecting subcortical structures. Furthermore, we maintained isoflurane concentrations below 2%, which was the level used by He et al. ^84^ It therefore is possible, albeit unlikely, that some degree of our observations reflected FUS modulation of state of anesthesia. Nevertheless, the absence of any significant overall diameter changes in sham experiments offers evidence that the post-FUS dilation did not reflect global physiological trends that evolved during the course of the experiment.

Ultimately, the broad concept of “persistent effects” owing to FUS neuromodulation likely reflects a conglomeration of neural, astrocytic, and vascular phenomena. We observed vascular dynamics in microvasculature that unexpectedly differs significantly from the responses in larger vessels that feed into the vascular network. Taken together with the recent *in vitro* findings that pericytes are FUS-sensitive, further *in vivo* experiments will be critical to parse the various underlying mechanisms and cellular origins that lead to short and longer-term neural and behavioral effects. It should also be noted that while the current report focused on morphological changes, future studies utilizing modified versions of apparatus may be able to quantitatively report on blood flow changes. Hwang et al., for example, recently demonstrated the ability to extract surrogate indicators of blood flow in capillaries using an OCT device that employed a swept-source OCT system^94^. Integrating flow-direction sensitive OCT-A could also help distinguish arterial vs. venous flow in the cerebrovascular networks. Given differences in their role in neurovascular coupling, we would expect to find differential effects. Additionally, integrating two-photon microscopy in-line with OCT angiography ^95^ could permit us to individually correlate Ca^2+^ dynamics, in various cell types, with localized microvascular blood flow. A more holistic understanding of all the mechanisms of FUS neuromodulation could also increase the predictive accuracy and relevance of computational modeling. For noninvasive electromagnetic modalities such as TMS or tDCS, direct effects on electrophysiological function can be simulated using standard finite element methods ^96–98^. An equally representative capability to do so for FUS would enable more systematic, hypothesis-driven experimental design.

## Methods

### Surgical procedures

All animal experiments were performed in accordance with the guidelines of the institutional Animal Care and Use Committee of New York Medical College. Adult C57BL/6J mice between two to five months were anesthetized with an initial dose of 5% isoflurane (vaporized in medical grade compressed oxygen) for < 20s and was later maintained at 1.5%. Following initial anesthesia induction, animals were positioned in a stereotaxic apparatus (Stoelting Co., Wood Dale, IL) and eyes were covered with petrolatum ophthalmic ointment (Puralube^®^, Fera Pharmaceuticals, Locust Valley, NY). Mice were administered buprenorphine (0.5mg/kg, subcutaneously) and dexamethasone (0.2mg/kg, subcutaneously) to reduce inflammation, and body temperature was measured and maintained at 37℃ with a closed-loop temperature-controlled heating system that included a heating pad (TC-1000, CWE, Ardmore, PA) and rectal probe (40-90-8D, FHC, Inc., Bowdoin, ME). Respiratory rate was monitored and maintained at ∼100 breaths/min during the surgical and experimental procedures.

In preparation for the craniotomy procedure, the scalp was infused with lidocaine delivered subcutaneously and a midline incision was performed on the region overlying the forelimb’s representation on the primary somatosensory cortex (at the same point on the rostrocaudal axis as bregma and ∼2.5 mm lateral). A custom-fabricated aluminum headbar was affixed to the left side of the exposed skull with cyanoacrylate adhesive (Loctite 454, HenKel, Wisconsin) to improve positioning for surgical and experimental procedures. A 2.5 × 3 mm rectangular craniotomy was made using a handheld dental drill (Osada Model EXL-M40), being careful not to damage the dura mater. To maximize combined optical and acoustic transparency, the exposed brain was covered with a 2.5 × 2.5 mm square piece of poly(4-methyl-1-pentene) (PMP) (Goodfellow Cambridge Ltd, Huntingdon, England) (Koekkoek et al., 2018). The gap between the outer edge of the PMP window and the inner edge of the craniotomy was filled with colorless surgical silicone adhesive (Kwik-Sil, World Precision Instruments, Florida), and the PMP window was sealed to the skull with instant adhesive gel (Loctite 454, HenKel, Wisconsin). The metal headbar and exposed skull were covered with dental cement (C&B Metabond, Parkell, New York). Animals were allowed to recover for four days before imaging and sonication experiments and euthanized directly following imaging sessions via cervical dislocation.

### Optical coherence tomography angiography (OCT-A) system and data acquisition

We built a customized Spectral Domain OCT (SD-OCT) shown in Fig. 1A as our OCT-A device. The system incorporates a superluminescent diode light source (Superlum, cBLMD-T-850-HP, *λ*_c_ = 850*nm*, Δ*λ* = 160*nm*) and a spectrometer (Cobra-S 800, Wasatch Photonics) with a line-scan camera (Teledyne e2v) operating at 80 kHz. A 5X objective lens (M Plan Apo NIR, Mitutoyo) with a long working distance was installed in the sample arm. The system’s axial and lateral resolution are ∼2.5 µm and ∼3.5 µm in tissue, respectively. As depicted in Fig. 1A, there was an air gap between the objective lens and entry window of the ring transducer; we compensated for this by adjusting the path length of the reference arm of the OCT system.

Custom OCT-system software written in C++ and CUDA was developed to control hardware, including laser control, galvanometers for scanning, and cameras for image acquisition. The OCT images were acquired and processed in near real-time with GPU processing on a high-performance graphic card (GeForce RTX 3090, NVIDIA). A CUDA-based accelerated DFT registration and Split Spectrum Amplitude Decorrelation algorithm ^99^ developed to produce OCT-A images. Angiography processing separated the static reflectance signal from the higher dynamic OCT signal that arises from flowing blood, resulting in relative blood flow images. Each 3D OCT anatomical volumetric snapshot contained 600 x 600 cross-sectional planes (A-scans × B-scans). repeated frames per B-scans over the same location were acquired for OCT-A. The total acquisition and GPU processing time for each angiographic volume was 30s. *En face* projections were generated from flattened OCT-A images using MATLAB (MathWorks, Natick, MA).

### In vivo sonication protocol

A total of 28 animals were used in experiments including 10 animals sonicated with 1 W/cm^2^, 10 animals sonicated with 10 W/cm^2^, 3 sham animals, and 5 animals used separately for immunohistochemical analysis only. Each animal was used in only one recording session. Animals were induced with 5% isoflurane, then maintained at 1-1.5%. Respiration was monitored continuously and anesthesia depth was assessed periodically via toe-pinch pedal reflex. As depicted in Fig. 3D, in each recording session, mice were sonicated twice with the same intensity to explore not only a single sonication effect, but to capture potentially cumulative effects of multiple doses. To minimize overlap of acute responses, repeated sonication doses were delivered with a 10-minute separation. This timing was informed by prior reports that acute hemodynamic responses largely subside within 10 seconds of sonication^56–58^. Experimental results from these experiments were included in analysis if the animal survived and maintained a stable plane of anesthesia and physiological state during full course of the recording session, which typically lasted 40 minutes from induction to completion. Sham experiments were performed on separate cohorts of animals that were subject to surgical procedures (including implantation, recovery periods, and anesthesia protocol) that were identical to animals used in experiments that involved sonication. This both OCT-A imaging experiments as well as the separate cohort used for immunohistochemical assessment. For functional sonication experiments, animals were situated under the ultrasound transducer prepared the same as it was for sonication experiments (i.e. filled with degassed water and coupled to the head with ultrasound gel), however sonication was not applied. OCT-A imaging was performed at the same exact time periods as those in experiments with sonicated animals. Statistical analysis was performed using GraphPad Prism-10 (GraphPad Software, CA, USA). Specific tests used for different analyses are described explicitly in the text.

### Combined optical and acoustic apparatus

To permit simultaneous FUS application and OCT-A angiographic portraits of blood flow, we used a custom mount that permitted fine, 3-axis positioning for a wide-aperture FUS ring transducer (customized H-205B, Sonic Concepts, Inc., Bothell, WA), described in ^100^. The FUS transducer’s fundamental frequency was 510 kHz and achieved a focal spot of lateral width ∼3.3 mm (full width at half-maximum) which effectively over-filled the field of view. For the OCT-A modality, the FUS transducer was integrated into the beam path ∼2 cm below the primary objective lens. Animals were positioned on a platform atop a lab jack for coarse adjustments (Thorlabs, Inc., NJ). Stimulus waveforms were generated and amplified by a TPO-106 transducer drive system (Sonic Concepts, Inc., Bothell, WA). Sonication was delivered as a 20 s burst 510 kHz ultrasound delivered as a series of 0.5 ms pulses at a repetition rate of 1 kHz (i.e. 50% duty cycle). Two intensities were used, a low of 1 W/cm^2^ and a high of 10W/cm^2^ (both *I_sppa_*). This yielded peak pressures of 0.124 MPa and 0.45 MPa, respectively. Beam properties were characterized in a tank of de-gassed water using a RESON spherically directional hydrophone.

To confirm the colocalized acoustic and optical foci, we used a thermochromic liquid-crystalline sheet (peak thermochromic range 30-35℃) to visualize the FUS focus centroid. Because the OCT illumination center wavelength was near-infrared, the scanned beam OCT beam could be resolved as a “crosshairs” on the thermochromic sheet to correct for any mismatch in position relative to the FUS focus.

### Image processing and analysis methods

Image stacks were first processed to remove background noise and co-register to account for any positioning drift between repeated scans. We utilized Top-hat filtering (6.7 μm window width) to segment blood vessels and bin by diameter, then applied functions from the Vessel Analysis plugin in ImageJ (Fiji)^101^ for arterioles (15 – 20 μm diameter) for binarization and skeletonization. Smaller vessel analysis utilized additional functions from the VasoMetrics toolbox, which has been verified for use in quantifying both OCT-A and two-photon microscopy data^102^. A visualization of the vessel segmentation analysis is depicted in Fig. S3, which also illustrates that vessel branches could be tagged and tracked individually throughout the course of FUS application. Given the OCT-A system’s resolution limitations, the minimum inclusion diameter of 5 μm. To mitigate the inherently lower signal-to-noise ratio for vessel branches of small diameter, the average diameter was calculated based on the average of a series of finely spaced, automatically selected cross sectional lines. Cross-sectional line spacing along vessel branches was typically 5 μm, or about 15 lines for the smallest vessel branches.

To best resolve the depth profile of vessels (Fig. 4), we axially binned image volumes into 25-μm divisions in depth and performed vessel segmentation within those narrow ranges. Because the surface of the brain is neither perfectly flat nor precisely perpendicular to the scanning beam, prior to this processing, image volumes were de-tilted using custom Matlab code. This helped ensure that our depth characterization of vasculature was as accurate as possible.

### Immunohistological procedures

Brains were dissected out and postfixed in 4% paraformaldehyde for 24h at 4 ℃, after which they were washed in phosphate buffered saline solution (PBS) for one hour then equilibrated in 30% sucrose. Brains were then embedded in OCT compound (Tissue-Tek, Sakura Finetechnical Co., Tokyo, Japan) and sectioned into 18 µm thick coronal sections on a cryotome. Sections were washed and blocked by incubation with 10% normal goat serum (NGS) in PBS supplemented with 0.4% Triton X-100 for 1h at room temperature. They were then incubated with CD31 Rat Monoclonal Antibody (PECAM-1) (1:500 dilution, Abcam 56299) in 2% NGS and 0.3% Triton X-100 at 4℃ overnight. Sections were then incubated for an hour with Alexa-488 labeled secondary antibody (1:500 dilution, goat anti-rat, Abcam150165), after which sections were mounted on slides, dehydrated, and coverslipped with non-fluorescent mounting medium (Krystalon, 64969-95, EMD, Darmstadt, Germany).

Blood-brain barrier disruption was visualized based on the leakage of intraperitoneally administered Evans blue dye into the brain’s parenchyma. Half an hour prior to FUS treatment, 0.1mL of a 1% solution of Evans blue dye was injected intraperitoneally. FUS was administered as described above for assessing changes in CD31 distribution. The brain was then dissected out and post-fixed, then prepared as described above for CD31 analysis.

Distribution of Evans blue dye and CD31 was observed using fluorescence imaging with an All-in-One Fluorescence Microscope (Keyence, BZ-X810, Osaka, Japan). The illumination and emission filter parameters for observing Evans blue were 560 ± 28 nm (excitation) and 645 ± 38 nm (emission); for CD31, filter parameters were 470 ± 40 nm excitation, 525 ± 50 nm emission).

## Data and Code Availability

The data that support the findings of this study as well as the code used for analysis are available on request from the corresponding author, J.A.N.F.

## Competing Interests

The authors declare no competing interests.

## Author Contributions

Y.S. performed experiments, analyzed data, and prepared the manuscript, J.V.J. and A.P.R. performed data analysis, J.H. performed experiments, M.A., C.R., S.H., and C.Z. engineered the experimental apparatus and contributed to preparing the manuscript, J.A.N.F. performed experiments, performed data analysis, and prepared the manuscript.

## Acknowledgements

The authors would like to acknowledge Furong Hu for assistance with immunohistological analysis, tissue preparation and related imaging, Aparna Singh and Elisa Konofagou for assistance calibrating ultrasound systems, and CheukHimIan Lam for assistance with image processing. This work was supported by award R01EB028319 from the National Institute for Biomedical Imaging and Bioengineering to J.A.N.F. as well as R01EB025209, R01HL156265 and R21EB03268401A1 to C.Z.

---

**Figure S1:**
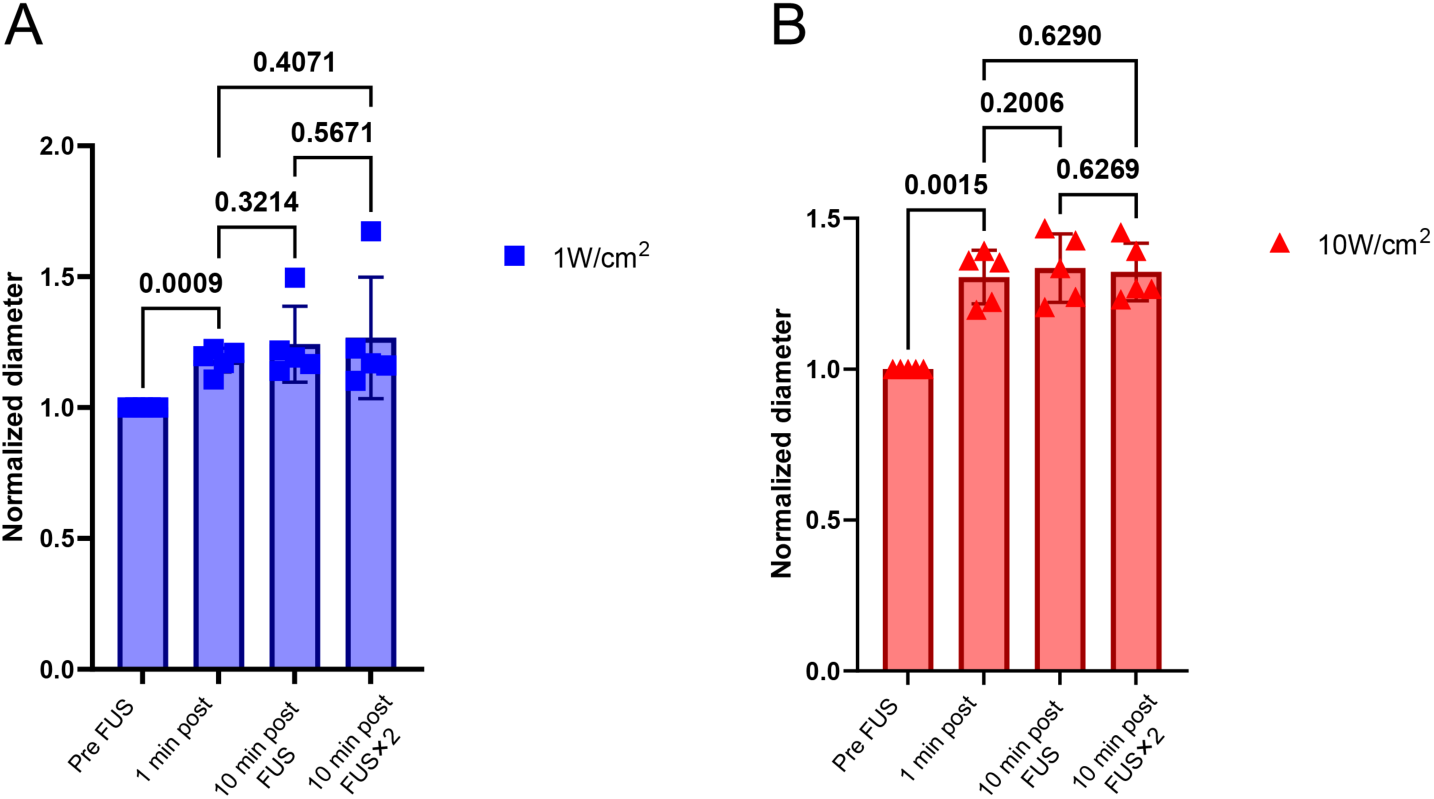
Diameter changes in response to repeated doses of FUS at (**A**) 1 W/cm^2^ and (**B**) 10 W/cm^2^ (*I_sppa_*). Individual data points correspond to results from individual animals (*N* = 10 for **A**, and a separate *N* = 10 for **B**). The data is derived from vessel branches of diameter < 15 μm. *P*-values were obtained via one-way ANOVA and Fisher’s Least Significant Difference test.

**Figure S2:**
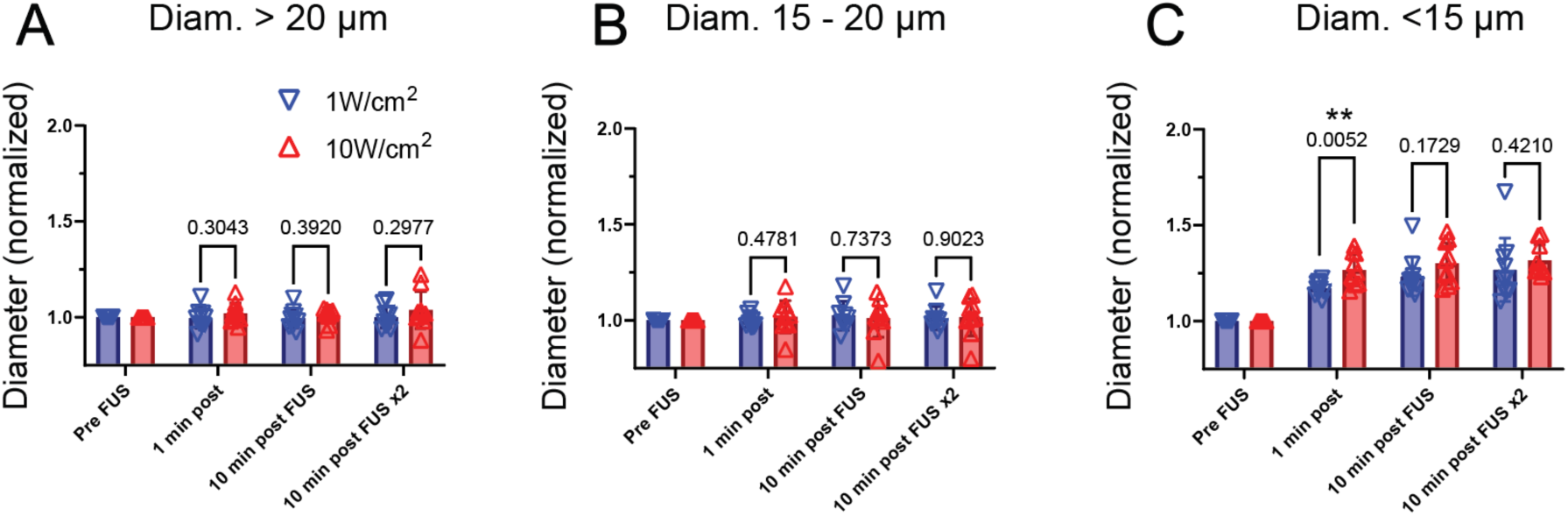
Sonication intensity dependent impact on vascular dilation. Panels **A**-**C** respectively depict normalized diameter changes in vascular branches of pre-FUS diameter > 20 μm, 15-20 μm, and < 15 μm in response to sonication at of 1 W/cm^2^ (*N* = 10) and 10 W/cm^2^ (*I_sppa_*) (separate *N* = 10). *P*-values listed above brackets are obtained via one-way ANOVA. For vessel branches of diameter < 15 μm, 1 W/cm^2^ and 10 W/cm^2^ evoked changes that were significantly different within the 1 minute directly following sonication, but not for repeated doses or later time periods.

**Figure S3:**
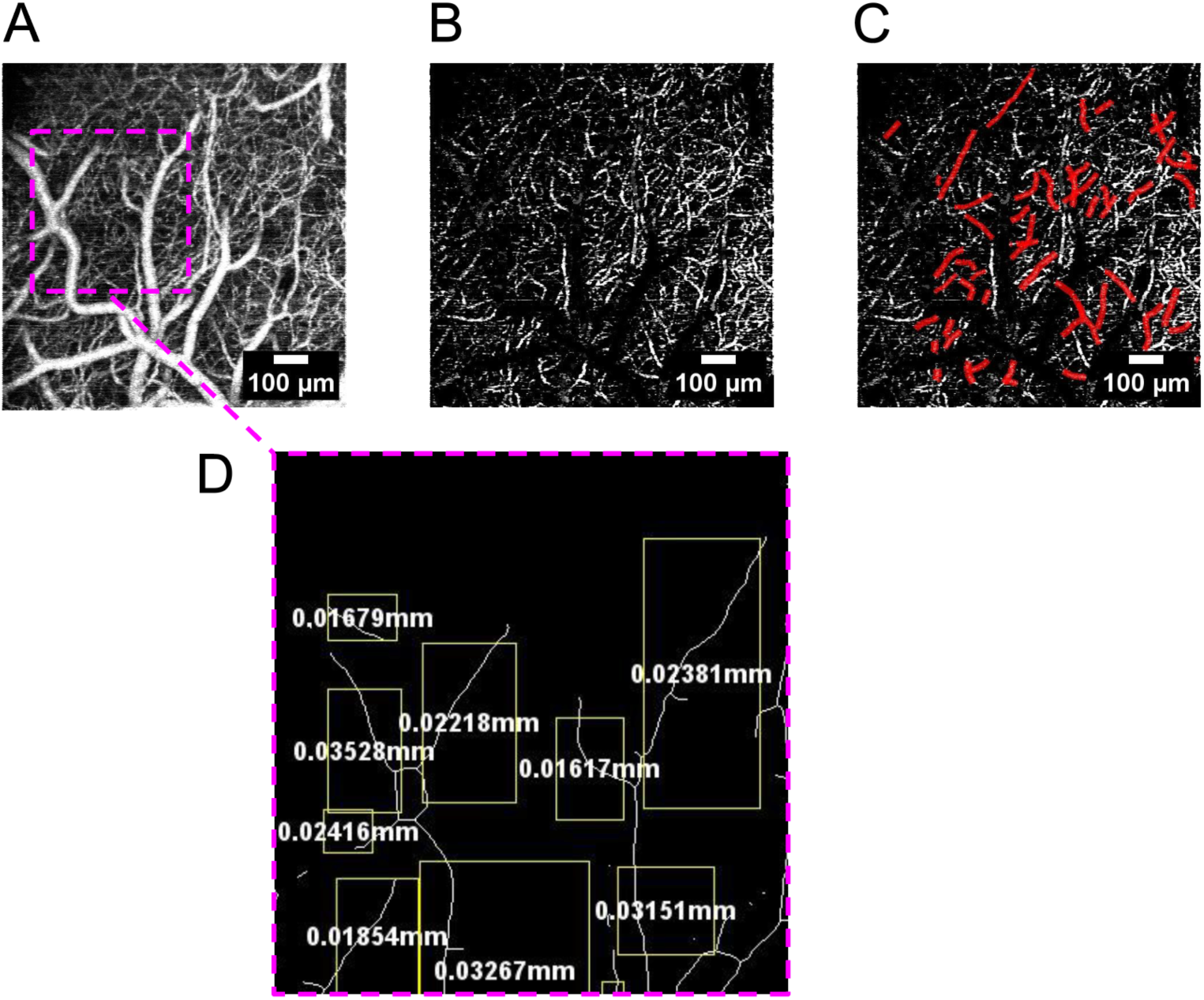
Diameter-specific analysis of angiographic measurements. **A** depicts a *z*-projection of a 1 mm × 1 mm × 540 μm region of somatosensory cortex *in vivo*. **B** illustrates diameter-specific segmentation of microvascular diameter vessel branches; red vessel branches in **C** were auto-selected for analysis because they could be resolved throughout all conditions. **D** illustrates, in a magnified subregion of **A**, the vessel “tagging” feature of our data analysis pipeline, which enabled us to track individual vessel branches through the course of FUS application.

**Figure S4:**
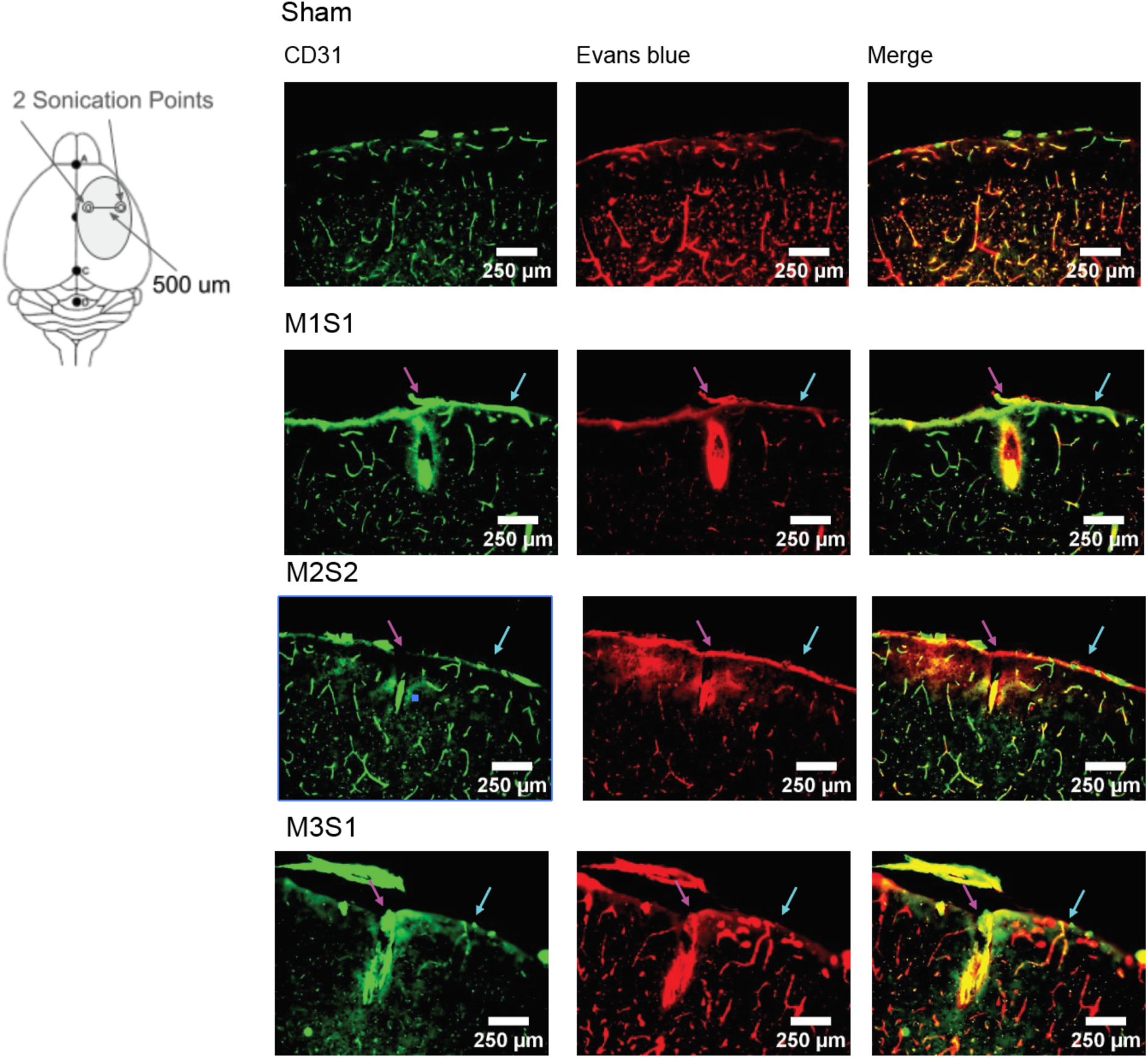
Results from additional immunohistochemical assessments of vascular integrity and BBB permeabilization. An illustration of the relative coordinates on the brain where FUS was targeted is shown at left. The more medial location received higher intensity sonication (>200 W/cm^2^ *I_sppa_*, 100% duty cycle, 20 s), while the lateral location was sonicated at a lower intensity (1.2 W/cm^2^, 100% duty cycle for 20s). Each row on the set of panels on the right show coronal sections obtained at the rostrocaudal axis location where sonication was performed for individual experiments (distinct from the results shown in Fig. 5). The magenta and cyan arrows represent, respectively, high-and low-intensity sonication locations. The top row shows results of a sham animal, which was subject to the same surgical procedures and mounting under transducer, yet no sonication was delivered.

**Figure S5:**
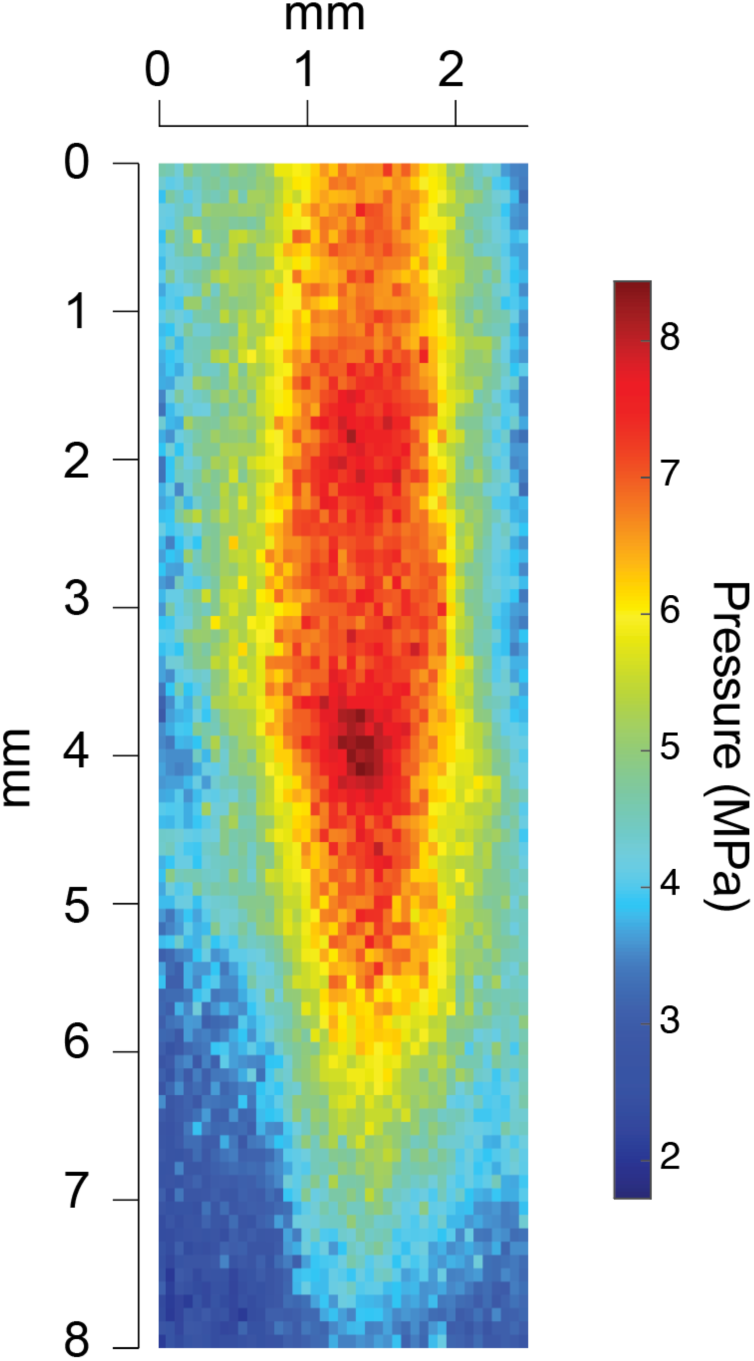
Axial pressure profile of the FUS beam, characterized in a tank of de-gassed water using a RESON spherically directional hydrophone. The horizontal axis corresponds to the plane parallel to the surface of the imaged surface, and the vertical axis corresponds to axial depth in the tissue. On the vertical scale, “0” corresponds to the bottom exit plane of the ring transducer, which is above the mouse’s head because the transducer’s focal curvature begins within the ring. The mouse’s skull was positioned at the beam’s actual focus.

## Notes

### Competing Interest Statement

The authors have declared no competing interest.

### Summary of Updates

Manuscript results updated to reflect new depth-resolved analysis; Figures 2, 3, 4, and 5 revised; new supplementary figures added; New author added for added expertise.

