## Supplemental figures for "Label free, capillary-scale blood flow mapping *in vivo* reveals that low intensity focused ultrasound evokes persistent dilation in cortical microvasculature"

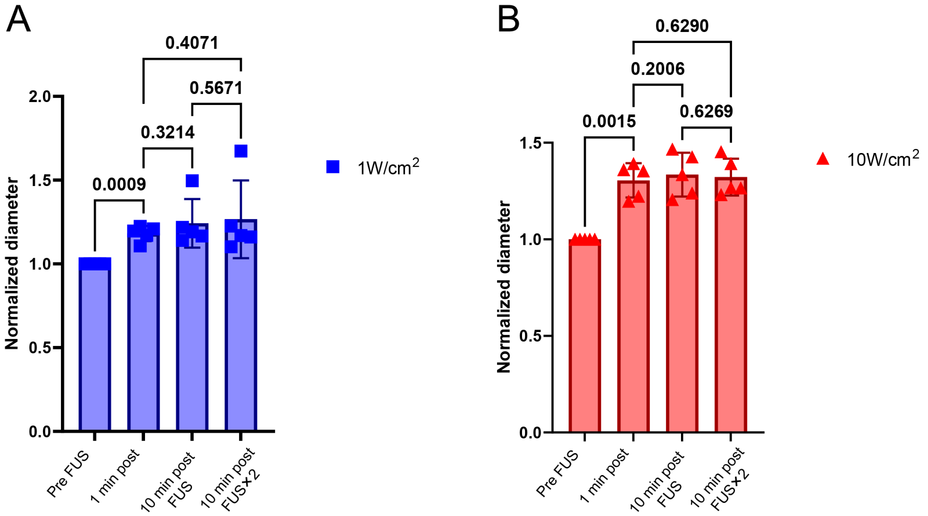


**Figure S1**: Diameter changes in response to repeated doses of FUS at (**A**) 1 W/cm$^2$ and (**B**) 10 W/cm$^2$ ($I_{sppa}$). Individual data points correspond to results from individual animals ($N$ = 10 for **A**, and a separate $N$ = 10 for **B**). The data is derived from vessel branches of diameter < 15 µm. *P*-values were obtained via one-way ANOVA and Fisher's Least Significant Difference test.

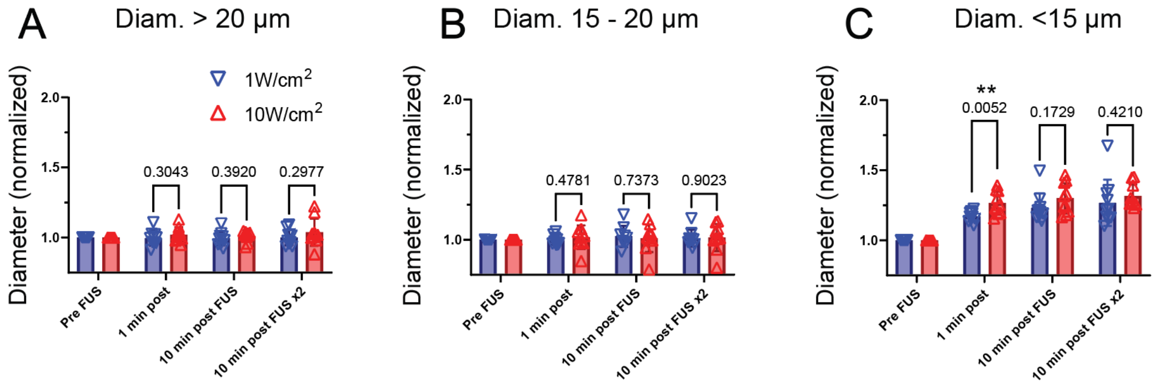


**Figure S2:** Sonication intensity dependent impact on vascular dilation. Panels **A**-**C** respectively depict normalized diameter changes in vascular branches of pre-FUS diameter > 20 µm, 15-20 µm, and < 15 µm in response to sonication at of 1 W/cm$^2$ ($N$ = 10) and 10 W/cm$^2$ ($I_{sppa}$) (separate $N$ = 10). $P$-values listed above brackets are obtained via one-way ANOVA. For vessel branches of diameter < 15 µm, 1 W/cm$^2$ and 10 W/cm$^2$ evoked changes that were significantly different within the 1 minute directly following sonication, but not for repeated doses or later time periods.

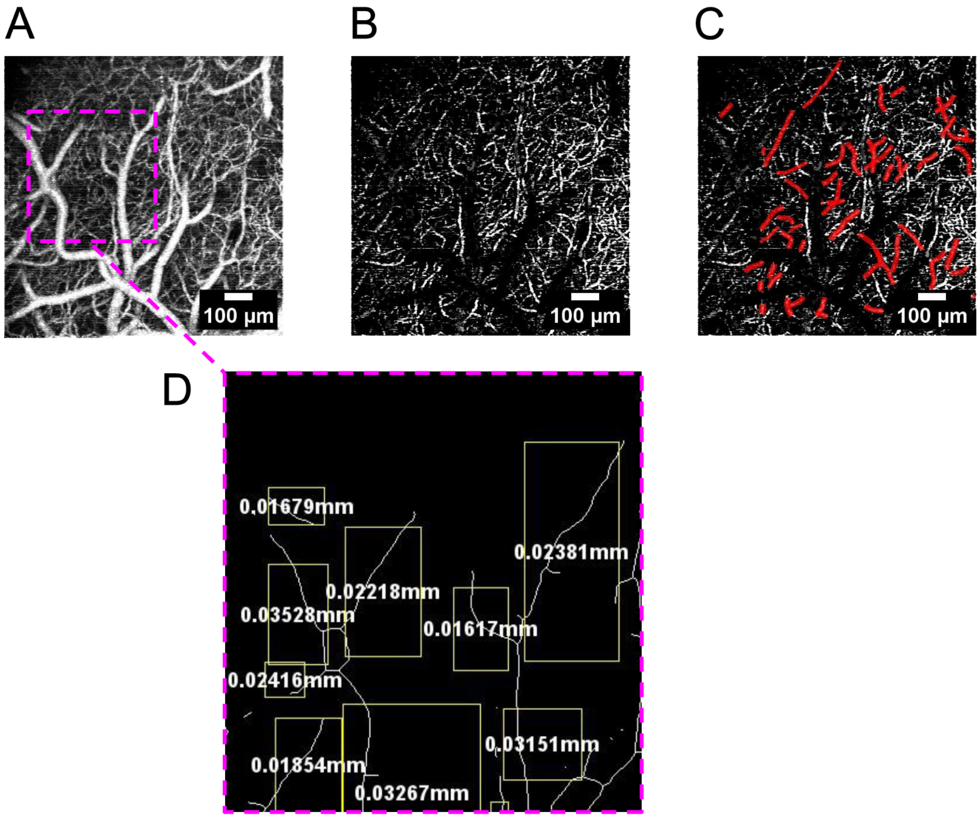


**Figure S3**: Diameter-specific analysis of angiographic measurements. **A** depicts a $z$-projection of a 1 mm × 1 mm × 540 μm region of somatosensory cortex *in vivo*. **B** illustrates diameter-specific segmentation of microvascular diameter vessel branches; red vessel branches in **C** were auto-selected for analysis because they could be resolved throughout all conditions. **D** illustrates, in a magnified subregion of **A**, the vessel "tagging" feature of our data analysis pipeline, which enabled us to track individual vessel branches through the course of FUS application.

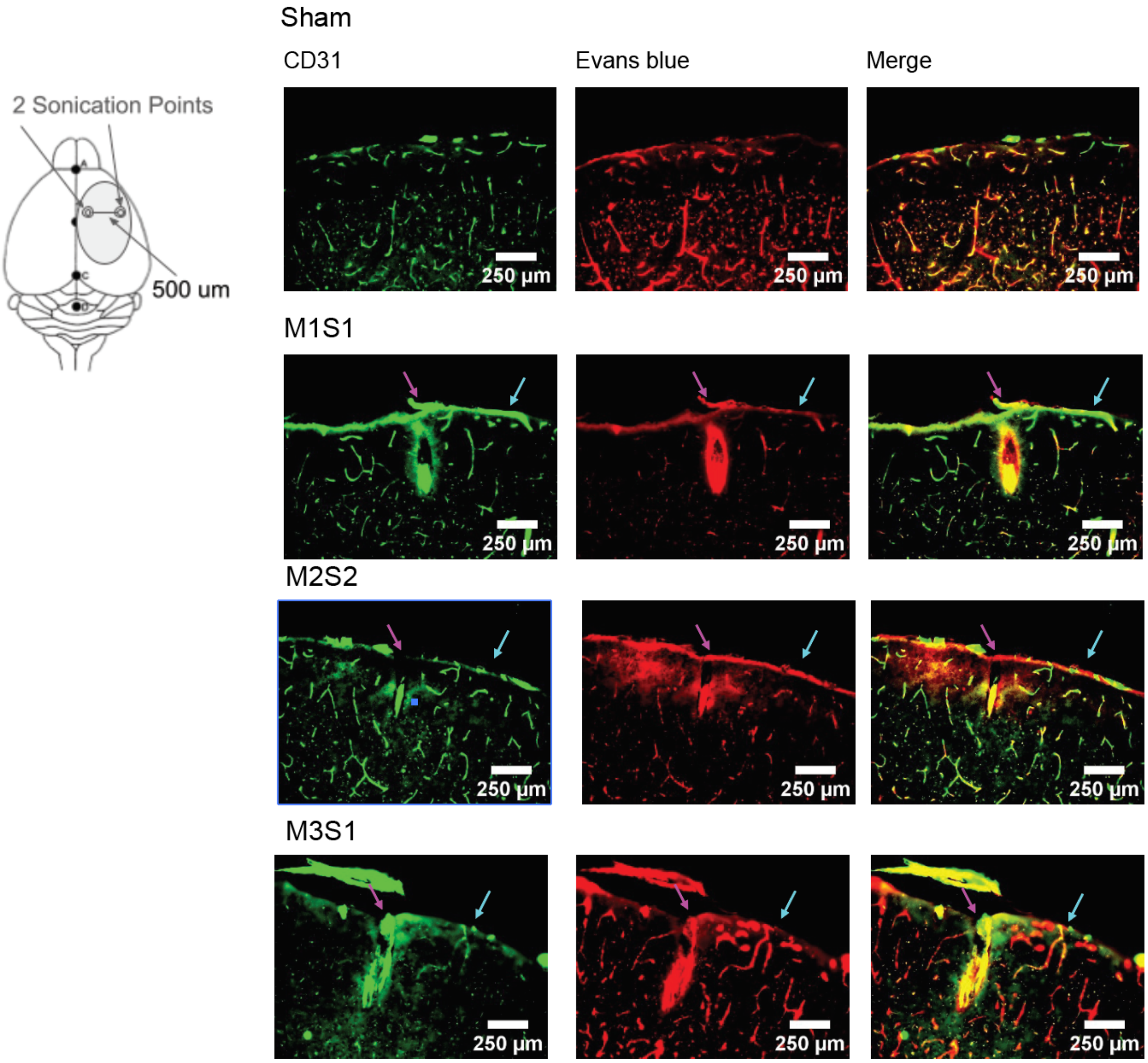


**Figure S4:** Results from additional immunohistochemical assessments of vascular integrity and BBB permeabilization. An illustration of the relative coordinates on the brain where FUS was targeted is shown at left. The more medial location received higher intensity sonication (>200 W/cm$^2$ $I_{sppa}$, 100% duty cycle, 20 s), while the lateral location was sonicated at a lower intensity (1.2 W/cm$^2$, 100% duty cycle for 20s). Each row on the set of panels on the right show coronal sections obtained at the rostrocaudal axis location where sonication was performed for individual experiments (distinct from the results shown in Fig. 5). The magenta and cyan arrows represent, respectively, high- and low-intensity sonication locations. The top row shows results of a sham animal, which was subject to the same surgical procedures and mounting under transducer, yet no sonication was delivered.

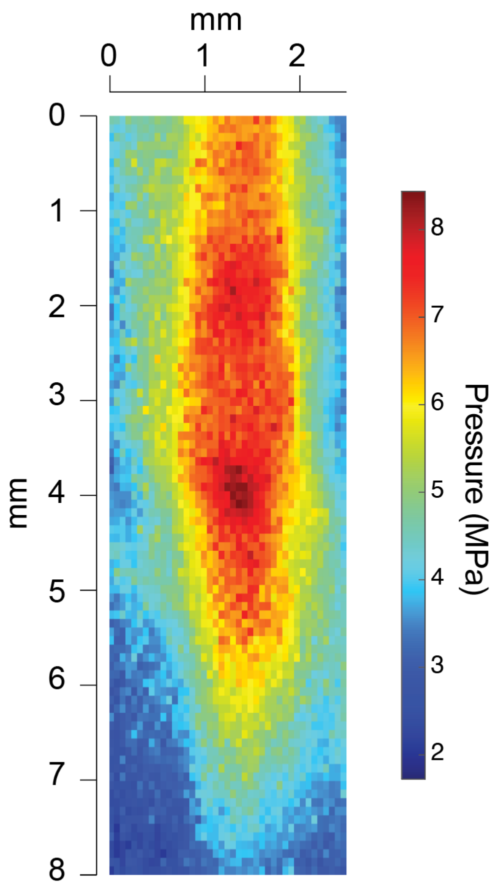


**Figure S5**: Axial pressure profile of the FUS beam, characterized in a tank of de-gassed water using a RESON spherically directional hydrophone. The horizontal axis corresponds to the plane parallel to the surface of the imaged surface, and the vertical axis corresponds to axial depth in the tissue. On the vertical scale, “0” corresponds to the bottom exit plane of the ring transducer, which is above the mouse’s head because the transducer’s focal curvature begins within the ring. The mouse’s skull was positioned at the beam’s actual focus.
